# Candidate proteins associated with popping expansion capacity of popcorn

**DOI:** 10.1101/2022.07.30.502124

**Authors:** Talita Mayara de Campos Jumes Gemelli, Isaac Romani, Natália Ferreira Dos Santos, Maria de Fátima P.S. Machado, Carlos Alberto Scapim, Gilberto Barbosa Domont, Fábio César Sousa Nogueira, Adriana Gonela

## Abstract

The mechanical resistance of the popcorn pericarp has a positive and direct relationship to its expansion volume. It allows enough time for the endosperm to gelatinize completely before its extravasation. Expansion is a polygenic trait that has been extensively studied. However, no records in the literature indicate proteins that directly affect pericarp thickness and integrity. Therefore, the present work aimed to identify candidate pericarp proteins associated with the expansion capacity of popcorn kernels using the shotgun proteomic approach. The analyses were carried out in the pericarp of two popcorn inbred lines, P11 (expansion volume of 30 mL g^-1^) and P16 (expansion volume of 14 mL g^-1^), in two developmental stages (15 and 25 DAP). A total of 803 non-redundant proteins were identified. Most of them were involved in key processes associated with pericarp development and thickening. Two candidate proteins stood out among the differentially abundant proteins. Peroxidase was up-accumulated in P11/25 DAP (high popping expansion) and was 1.498 times more abundant in this inbred line, while xyloglucan endotransglucosylase/hydrolase was more abundant in P16 (low popping expansion) in both developmental stages. Thus, the peroxidase protein increases expandability, whereas xyloglucan endotransglucosylase/hydrolase decreases it, even though its specific role has not been elucidated. These proteins should be further investigated, as they may be used to improve expansion capacity in popcorn breeding programs.

## Introduction

Popping is a physicochemical process in which the gelatinization and expansion of starch occur simultaneously through the rupture of the pericarp when grains are subjected to high temperatures (177-185 °C) for a short period [1, 2]. In this context, popcorn maize [*Zea mays* L. var. everta (Sturtev) L. h. Bailey] is the species that presents the best results to this process, as it has grains with morphological, physical-chemical, and genetic characteristics that favor expansion [1, 3, 4, 5].

The pericarp of popcorn kernels performs one of the most important functions during popping. It is more structurally organized in popcorn than in common corn [6], maintaining the integrity of the kernel. Furthermore, it provides resistance to the increase in internal pressure, reaching 760-930.8 kPa (or 10 atm). Finally, it increases heat-transfer efficiency into the kernel for an ideal period until starch gelatinization occurs [1, 6, 7]. Pre-existing ruptures in the pericarp cell layers decrease the kernel resistance to the pressure generated during heating, leading to internal steam escaping and producing smaller or even no pieces of popped corn [6].

In the species *Zea mays*, the pericarp is formed by cells with a type II primary cell wall, which stands out for having glucuronoarabinoxylan as the most abundant hemicellulose (40-50%) linked to cellulose microfibrils. These components form a complex network that confers rigidity and extensibility to the cell wall [8]. In turn, xyloglucan appears in lower abundance (1-5%); therefore, it is a secondary component [9]. However, even though it is less abundant, xyloglucan metabolism plays a more central role in restructuring the cell wall of commelinoid monocots, such as maize [10]. One of the main enzymes that promote the cleavage and reassembly of xyloglucan molecules is xyloglucan endotransglucosylase/hydrolase, which acts on β-1,4 or β-1,3 glycosidic bonds [11].

Lignin is the second most abundant component in pericarp cell walls, which provides hydrophobicity and mechanical strength [12], in addition to protection against biotic [13] and abiotic stresses [14]. Lignin results from the oxidative polymerization of three canonical phenolic alcohols [sinapyl (S unit), coniferyl (G unit), and ρ-coumaryl (H unit) alcohols] carried out by peroxidases and laccase enzymes in the secondary cell wall [14–17]. Peroxidases have specialized activities according to the organ, cell, and developmental stage. In conclusion, not only spatiotemporal regulation of gene expression and protein distribution, but also differentiated oxidation properties of each Prx define the function of this class of peroxidases [14].

Regarding the expansion capacity of popcorn, it is suggested that both lignin content and composition contribute positively to the expansion volume [18]. However, which genes/proteins are associated with lignin’s highest content and composition has not been determined. Thus, the present work aimed to identify candidate pericarp proteins associated with the expansion capacity of popcorn kernels through shotgun proteomics using iTRAQ^®^ (Isobaric Tags for Relative and Absolute Quantitation - Sciex).

## Materials and methods

### Popcorn inbred lines

Two inbred lines of popcorn (early cycle) with different expansion volumes, P11 (expansion volume 30 mL g^-1^) and P16 (expansion volume 14 mL g^-1^), were used in the quantitative proteome analysis.

The experiment was performed in the field (23°25’S, 51°57’W, 550 m altitude) using the complete block design. Plots contained four 5-m long lines spaced 0.90 m apart and plants spaced 0.20 m apart, with 10 repetitions. Irrigation was performed.

The plants were manually self-fertilized, and the ears were collected 15 and 25 days after pollination (DAP) (S1 Fig). Three ears of each treatment (lineages x developmental stage x repetition) were collected, totaling four treatments: T1 (P11/15 DAP), T2 (P16/15 DAP), T3 (P11/25 DAP), and T4 (P16/25 DAP). First, the grains were excised from the ears, and then the pericarps were removed, immediately frozen in liquid nitrogen, and stored in an ultra-freezer (−80 °C) (S1 Fig 1).

### Extraction and preparation of protein samples

After extracting proteins from the pericarp/aleurone [19], the samples were lyophilized and stored in an ultra-freezer (−80 °C). The protein concentration was obtained through the fluorimetric method using the Qubit^®^ 2.0 Fluorometer (Invitrogen). Afterward, the samples were diluted (1:10) in 200 mM TEAB (triethylammonium bicarbonate) buffer, and they were reduced with 10 mM DTT (dithiothreitol) for 1 hour at 25 °C. Subsequently, the samples were alkylated with 40 mM iodoacetamide (IAA) at room temperature in the dark.

Proteins (50 μg) were digested with trypsin (Promega, Madison, WI, USA) at a protein:trypsin ratio of 50:1. The samples were cleaned and purified using C-18 Macro SpinColumns^™^ (Harvard Apparatus). Peptide samples were resuspended in 50 mM TEAB buffer and quantified by the fluorimetric method using the Qubit^®^ 2.0 Fluorometer. Subsequently, each treatment was divided into 25 μg aliquots of peptides.

### iTRAQ labeling and SCX (cation exchange fractionation)

The peptides were processed according to the iTRAQ^®^ 4-plex protocol. Peptides from treatments T1 (P11/15 DAP), T2 (P16/15 DAP), T3 (P11/25 DAP), and T4 (P16/25 DAP) were labeled with reporter ions of 114, 115, 116, and 117 Da, respectively. Biological triplicates (G1, G2, and G3) were performed per treatment.

Peptide samples tagged with iTRAQ^®^ were resuspended in 100 μL of buffer A [10 mM KH_2_PO_4_, 25% acetonitrile (ACN), pH 3] and loaded onto a cation exchange Macro SpinColumn^™^ (Harvard Apparatus). The peptides were eluted at a gradient of buffer A for 10 minutes at room temperature, with subsequent centrifugation at 100 rpm for 3 minutes. This procedure was repeated twice, and the final eluate collected was transferred to a new tube and identified as flow-through (FT).

The first fractionation was performed by adding 300 μL of buffer B (10 mM KH_2_PO_4_, 25% ACN, 100 mM KCl) to the Macro SpinColumn. After centrifugation at 100 rpm for 3 minutes, the content was collected and stored in a new tube designated 100. Then, two more fractionation processes were carried out using 250 mM and 500 mM KCl, thus obtaining fractions of 250 and 500. Residual salts were removed from the cationic process using C-18 Macro SpinColumns^™^ (Harvard Apparatus).

### LC–ESI–MS/MS analysis based on LTQ Orbitrap

Peptide samples were solubilized in 0.1% formic acid and fractionated using a nano-LC Easy 1000 system (Thermo Fisher) coupled to an Orbitrap-type mass spectrometer (Q Exactive Plus, Thermo Scientific).

For each sample, 2 μg of peptides was applied to a trap column (200-μm inner diameter and 2-cm length) packed in-house with Reprosil-Pur C18 5-μm resin (Dr. Maisch^®^) (200 Å pores). The peptides were eluted in an analytical column (75-μm inner diameter and 18-cm) packed in-house with Reprosil-Gold C18 3-μm resin (Dr. Maisch^®^) (300 Å pores). Peptide separation was performed using a gradient from 95% solvent A (5% ACN and 0.1% formic acid) to 40% solvent B (95% ACN and 0.1% formic acid) over 120 minutes.

The Orbitrap mass spectrometer was controlled by Xcalibur 2.2 software, which was programmed to operate in automatic data-dependent (DDA) mode. The mass spectrum was acquired with a resolution of 70,000 to 200 m/z (mass/charge). The reading spectrum comprised peptides with 375 to 2000 m/z.

The 15 most intense ions were fragmented and then subjected to MS/MS acquisition using higher energy collision-induced dissociation (HCD). Each DDA consisted of a scan survey comprising a range of 200-2000 m/z. Peptides with undetermined charges and +1 were rejected. A 5% ammonia solution contained in a 15 mL tube with the lid open was placed close to the nESI region to avoid the effect of increasing ionic charge caused by the iTRAQ 4-plex [20, 21].

The experiment consisted of a total of 36 mass spectrometer runs, with samples derived from pericarp/aleurone, biological triplicates (G1, G2, and G3), four fractionation steps (100, 250, 500, and FT), and technical triplicates (runs in the mass spectrometer). All biological replicates and their respective fractionations were analyzed together.

### Data analysis

Raw files were visualized using Xcalibur v.2.1 software (Thermo Scientific), and data were processed using Proteome Discoverer v.1.4 software. *Zea mays* L. cv. B73 was obtained from the UniProt Consortium (http://www.uniprot.org/) (UniProt, 2019), which had 85,525 entries (downloaded in April 2016). Analyses were performed according to the following parameters: 10 ppm tolerance for precursor ion masses, MS accuracy of 0.1 Da for HCD, two missed cleavages allowed for fully tryptic peptides, carbamidomethylation of cysteine as a fixed modification, oxidation of methionine and lysine and N-terminal oxidation caused by iTRAQ^®^ as variable modifications.

Protein number and groups, peptide number, and the quantitative values of each marker were estimated using the Proteome Discovery software through the SEQUEST algorithm. The false discovery rate (FDR) was set at 1% for the detection of peptides and proteins. The proteins received the UniProt Consortium identification codes.

PatternLab for Proteomics software (http://www.patternlabforproteomics.org/) [22] was also used to identify the peptides and proteins. Briefly, the search was performed with the Comet search tool limited to fully tryptic candidate peptides, while cysteine carbamidomethylation and iTRAQ-4 (N-terminal and K) were defined as fixed modifications. A cut-off point of 1% was established for FDR at the peptide level based on the number of labeled decoys. This procedure was performed independently for each data subset, resulting in an FDR value independent of the tryptic or charge state. In addition, a peptide with a minimum length of six amino acids was required.

The results were post-processed to accept only PSMs with less than 6 ppm of the global identification average. One-hit wonders (proteins identified with only one mass spectrum) were considered only if the XCorr values were greater than 2.5. These criteria led the values of FDRs (at the protein level) to be less than 1% for all the results surveyed.

### Quantitative analysis of total proteins

The quantitative analysis for protein expression was conducted with PatternLab for Proteomics software using the Isobaric Analyzer module [22]. In this module, iTRAQ reporter ion intensities were extracted, applied to purity correction (as indicated in the manufacturer’s instructions), and then normalized by the signal from each isobaric marker according to the total ion current for that respective reporter ion mass. PatternLab provides features in its platform. Thus, we were able to perform a comparative data investigation based on ratios provided by the two experimental conditions.

In this study, analyses were performed based on the treatment ratios, with the results for lines P11 and P16 established in the numerator and denominator, respectively, for the two developmental stages (15 and 25 DAP). Thus, the ratios analyzed were RI (P11_15DAP_/P16_15DAP_) and RII (P11_25DAP_/P16_25DAP_).

The Isobaric Analyzer module employs a peptide-centric approach that assigns a paired t-test with p-values to each peptide and then converges to a final p-value (at the protein level) through Stouffer’s method. To conduct statistical analysis, we assumed log2-values of the integration value of the precursor (ICPL) or the reporter ion (iTRAQ) and normalized them on sets of three replicates. An overall average with all values and stages was estimated. By that, the samples became comparable by subtracting this average. Finally, we merged the values of all replicates (considering each developmental stage individually), and the total average was used to normalize each replicate. A protein’s log fold change was estimated by averaging the corresponding peptide log folds. Our differential proteomic comparison considered only proteins identified with unique peptides (i.e., peptides that map to a single sequence in the database), a paired t-test ≤ 0.05, and an absolute peptide fold change cut-off higher or less than 1.5x.

### Bioinformatics analysis

The proteins identified in the UniProt Consortium also received the official nomenclature for the species *Zea mays* available in MaizeGDB (Maize Genetics and Genomics Database) (https://www.maizegdb.org/).

The Search & Color pathways feature of KEGG Mapper (Kyoto Encyclopedia of Genes and Genomes) (https://www.genome.jp/kegg/mapper/convert_id.html) [23] together with MaizeMine v. 1.3 from MaizeGDB (http://maizemine.rnet.missouri.edu:8080/maizemine/templates.do) [24] were used to identify metabolic pathways. In addition, the proteins were subjected to functional categorization of Gene Ontology (GO) terms for biological processes with the aid of the Analysis Toolkit and Database for Agricultural Community program (AgriGO) (http://bioinfo.cau.edu.cn/agriGO/) version 2.0 [24]. The statistical parameters used in AgriGO were Fisher’s exact test and the multi-test adjustment method of Yekutieli (FDR under dependency) with a 5% significance level.

## Results

### Identified proteins

The protein and one-dimensional profiles obtained from the pericarp samples of the two popcorn lines (P11 and P16) in the four analyzed treatments, T1 (P11/15 DAP), T2 (P16/15 DAP), T3 (P11/25 DAP), and T4 (P16/25 DAP), showed a great abundance of proteins between 20-97 kDa, in addition to good reproducibility of the profiles (S2 Fig).

The quantitative characterization of the proteome based on iTRAQ^®^ identified 3,924 proteins that assumed an FDR of 1% in the samples of the two popcorn lines, thus determining high confidence in their identification. Of these, 803 were non-redundant and were analyzed in more detail (S1 Table). The 803 proteins were categorized into nine major biological processes (GO:0008150), eight molecular functions (GO:0003674), and seven cellular components (GO:0005575) (Fig 1, S2 Table).

**Fig 1.**
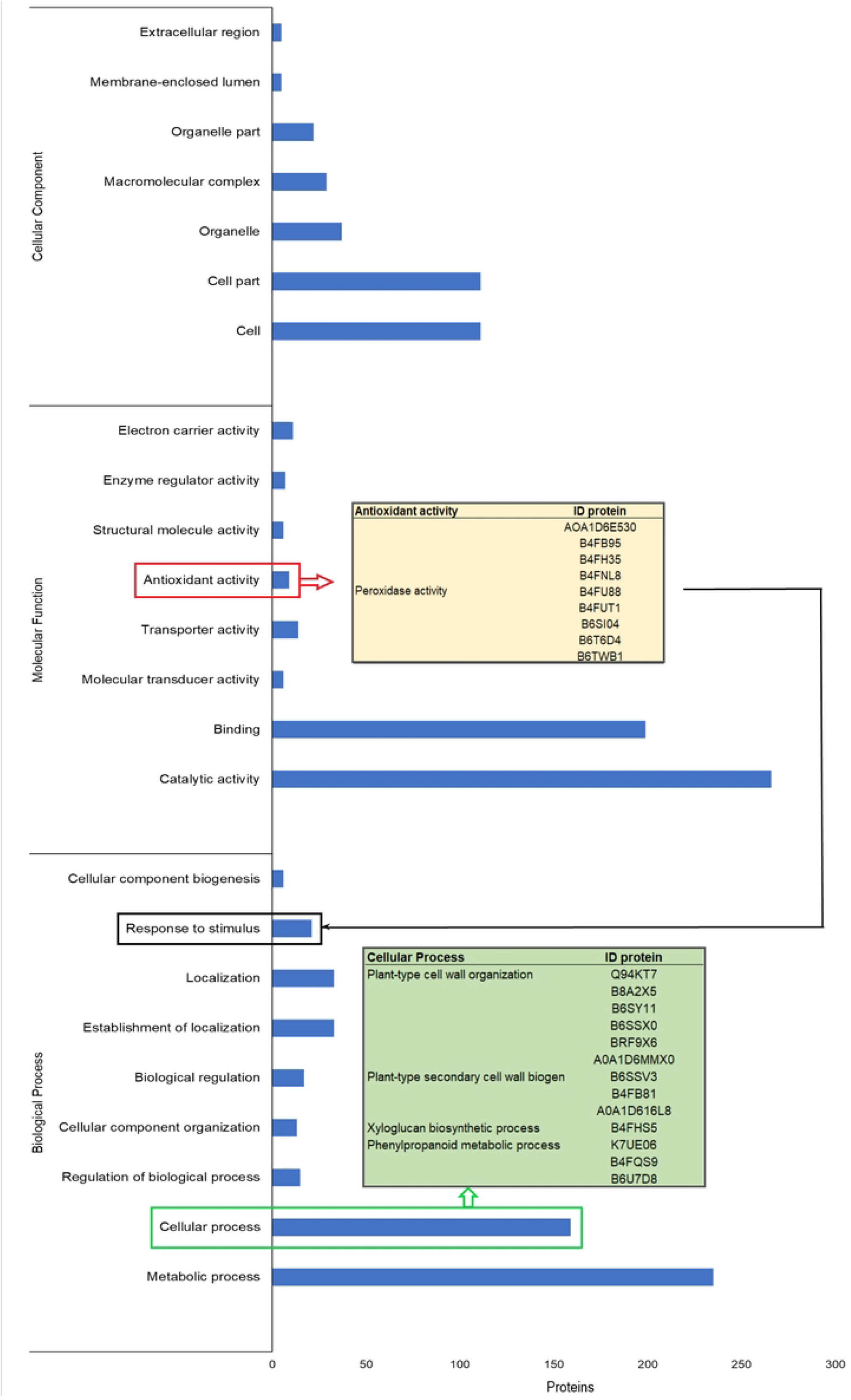
Main GO terms related to Biological Processes, Molecular Function, and Cellular Component of non-redundant proteins identified in the pericarp of popcorn inbred lines P11 and P16.

The categories that stood out the most in biological processes were metabolic (235 proteins) and cellular processes (159 proteins). Within the cellular processes, five proteins (Q94KT7, expansin; B8A2X5 and B6SSX0, pectinesterase; B6SY11, glycosyltransferase; B4F9X6, pectin acetylesterase, and A0A1D6MMX0, glycosyltransferase family 61 protein) were identified in the plant-type cell wall organization (GO:0009664); three proteins (B4FB81, fasciclin-like arabinogalactan protein 6; B6SSV3, fasciclin-like arabinogalactan protein 7, and A0A1D6I6L8, fasciclin-like arabinogalactan protein 8) were identified in plant-type secondary cell wall biogenesis (GO:0009834); two proteins (B4FQS9, trans-cinnamate 4-monooxygenase and B6U7D8, cinnamyl alcohol dehydrogenase) were linked to the lignin metabolic process (GO:0009808), and one protein (B4FHS5, xyloglucan endotransglucosylase/hydrolase) acted in the xyloglucan biosynthetic process (GO:0010411), as well as in cell wall biogenesis (GO:0042546) and organization (GO:0071555).

In the molecular function category, proteins were mainly allocated to catalytic activity (266 proteins) and binding (199 proteins). In addition, another category that deserves to be highlighted is antioxidant activity, in which nine proteins were assigned. All these proteins are peroxidases (A0A1D6E530, B4FB95, B4FH35, B4FNL8, B4FU88, B4FUT1, B6SI04, B6T6D4, and B6TWB1) with heme-binding activity (GO:0020037), metal ion binding (GO:0046872), and peroxidase activity (GO:0004601). Additionally, they are associated with response to stimulus.

### Differentially abundant proteins

Thirty-nine differentially abundant proteins were identified in the pericarp/aleurone of the two popcorn lines (Fig 2A, S3 Table). Early in development (15 DAP), eight differentially abundant proteins were identified (Fig 2A). Two proteins were up-accumulated exclusively in the inbred line P16 (low expansion capacity), K7UBG7 (cullin-associated NEDD8-dissociated protein 1) and B6T6D4 (peroxidase) (Figs 2A and 2B). These proteins were allocated to genetic information processing and phenylpropanoid biosynthesis pathways, respectively (Fig 3, S4 Table). Notably, the peroxidase B6T6D4 was 0.565 times more abundant in P16 (15 DAP) than in P11 (Table 1). In addition, three up-accumulated proteins were identified in P11 (high expansion capacity) at 15 DAP (Fig 2A). Two proteins, B6TI56 (ribose-5-phosphate isomerase) and B6ST80 (auxin-repressed 12.5 kDa protein), were up-accumulated exclusively at this developmental stage (Figs 2A and 2B), acting on the pentose phosphate and metabolic pathways, respectively (Fig 3, S4 Table).

**Fig 2.**
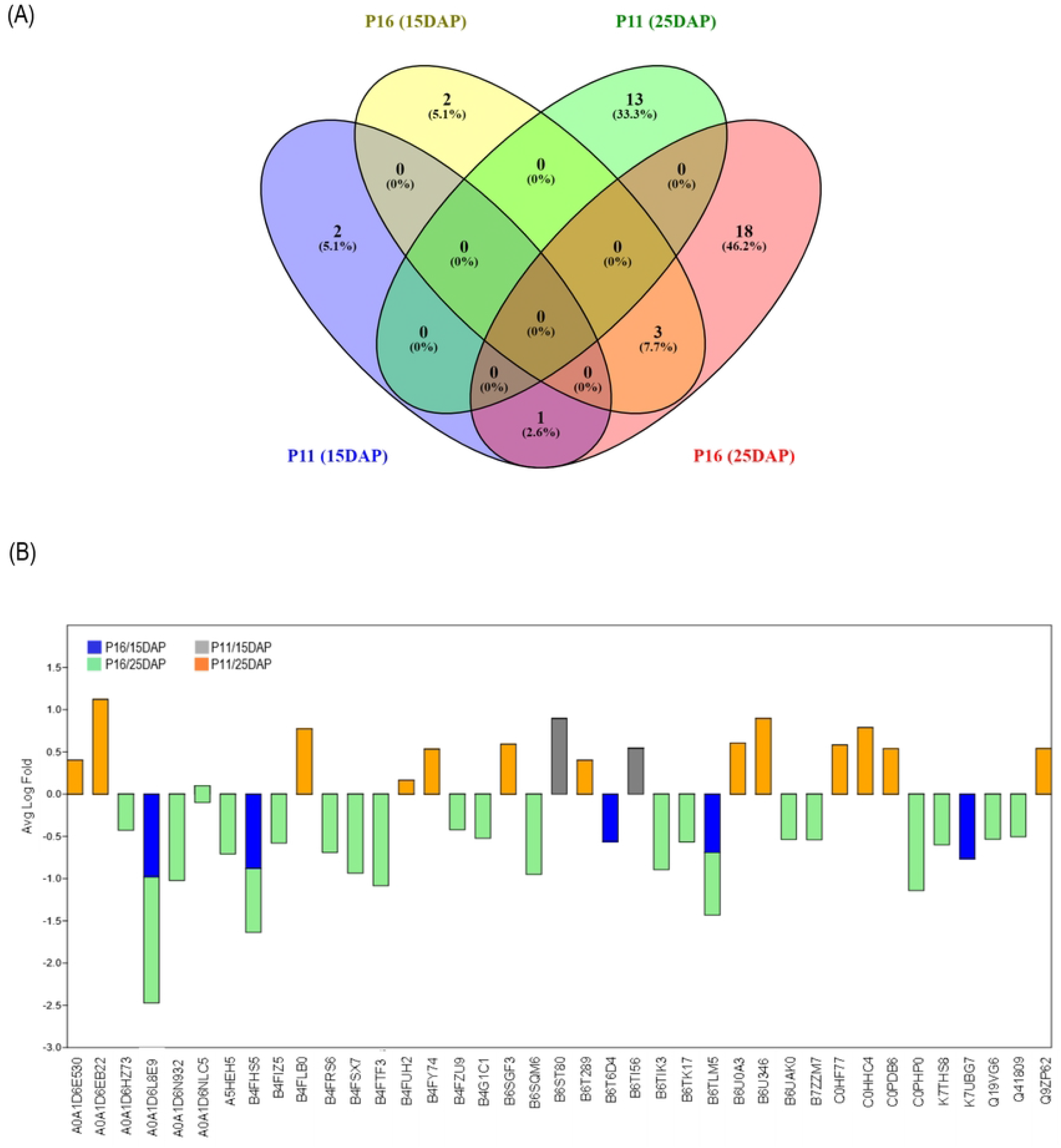
Differentially abundant proteins identified in two popcorn inbred lines (P11 and P16) in two developmental stages (15 and 25 DAP) (A); relationship of up-accumulated proteins in P16/15 DAP, P16/25 DAP, P11/15 DAP, and P11/25 DAP (B).

**Fig 3.**
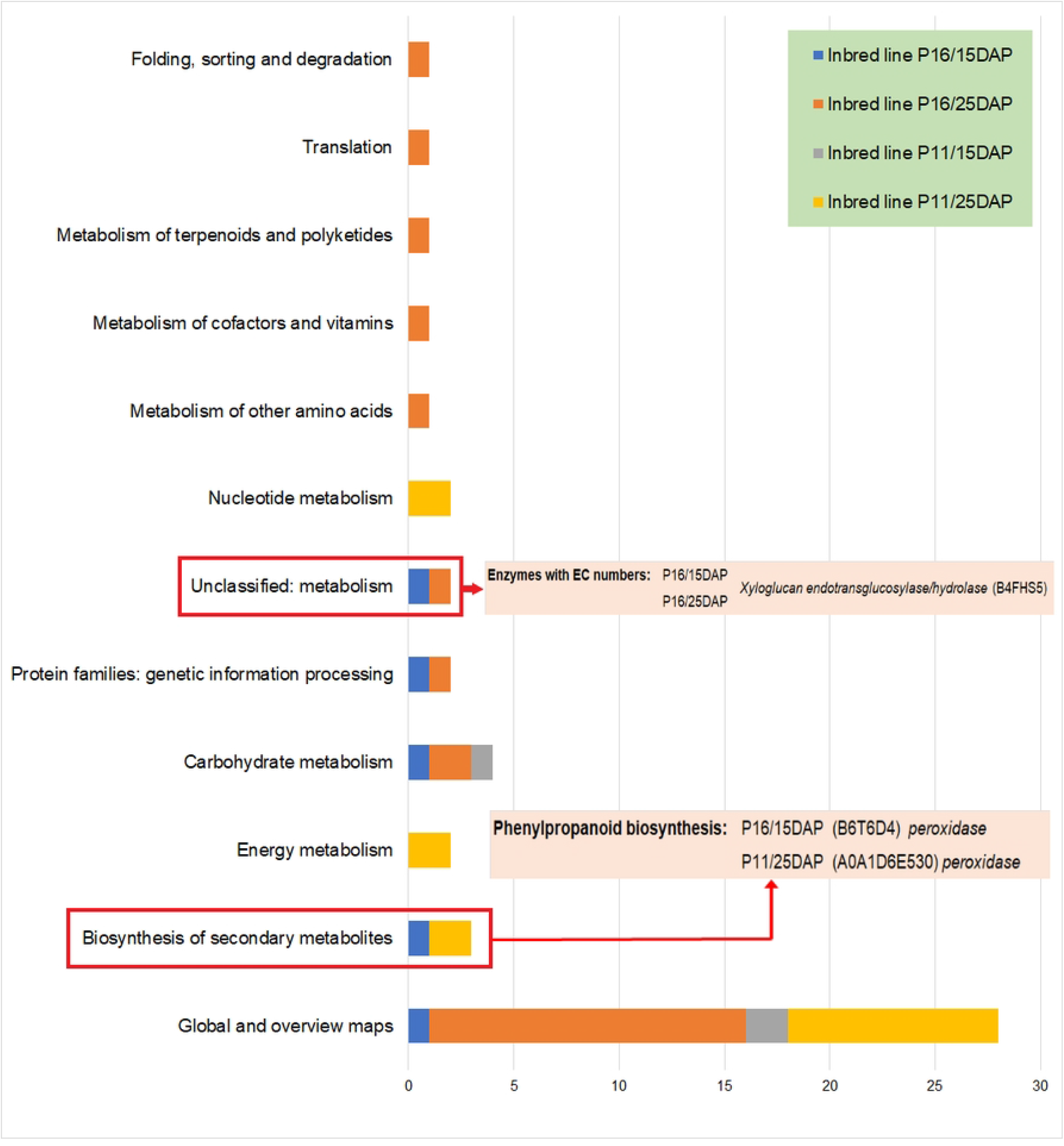
Metabolic pathways of differentially abundant proteins identified in the pericarp of two popcorn inbred lines (P11 and P16) in two developmental stages (15 and 25 DAP).

**Table 1.** Up-accumulated proteins identified in lignin and hemicellulose metabolic pathways.

| (Prot ID) Description | Log<sub>e</sub> (x) of ratios* |  | *p*-value |
| --- | --- | --- | --- |
|  | RI | RII |  |
| (B4FHS5) Xyloglucan endotransglucosylase/hydrolase | -0.881 |  | 0.010 |
|  |  | -0.756 | 0.010 |
| (B6T6D4) Peroxidase | -0.565 |  | 0.045 |
| (A0A1D6E530) Peroxidase |  | 0.404 | 0.010 |

\*Note: RI, 15 DAP (P11/P16); RII, 25 DAP (P11/P16).

At 25 DAP, among the 31 differentially abundant proteins identified, 18 were up-accumulated exclusively in P16 (low expansion capacity), and 13 were only upregulated in P11 (high expansion capacity) (Figs 2A and 2B, S3 Table). The vast majority of proteins, both in the P16 (13 proteins) and P11 (11 proteins) inbred lines, were identified in the global and overview map pathways (Fig 3). However, the up-accumulated proteins B4FZU9 (dihydropyrimidine dehydrogenase NADP^+^ chloroplastic), B4FSX7 (-+-neomenthol dehydrogenase), and B7ZZM7 (eukaryotic translation initiation factor 2 subunit alpha) were exclusively detected in P16. These proteins are related to the metabolism of other amino acids and metabolism of cofactors and vitamins, metabolism of terpenoids and polyketides, translation and folding, and sorting and degradation pathways. In turn, in P11 at 25 DAP, two proteins stood out, A0A1D6E530 (peroxidase) and B4FUH2 (aspartate aminotransferase), which were identified in the phenylpropanoid biosynthesis and amino acid metabolism pathways, respectively (Fig 3, S4 Table). Peroxidase was 1.498 times more abundant in P11 than in P16 at 25 DAP (Table 1).

In addition, xyloglucan endotransglucosylase/hydrolase (B4FHS5) stood out. This protein was up-accumulated exclusively in the inbred line P16 in the two stages of pericarp development (Fig 2B, S3 Table), and it was identified in the metabolism pathway (Fig 3, S4 Table). This protein was 0.414 and 0.469 times more abundant in the low expansion capacity line than in the high expansion capacity inbred line at 15 and 25 DAP, respectively (Table 1).

## Discussion

The popcorn pericarp is directly related to the expansion capacity. It has a mechanical resistance that allows it to withstand an internal pressure that can reach 930.8 kPa [1,6], thus allowing sufficient time for the endosperm to gelatinize completely before its rupture.

The pericarp thickness is directly associated with its mechanical resistance. Some studies have evaluated pericarp thickness [25,26], its type of inheritance [27] and integrity [7], and the role the cellulose matrix plays in the retention of moisture inside the seed [28] to correlate these traits with expandability. However, no work has sought to analyze the pericarp proteome to identify proteins associated with expansion capacity.

In the present work, most of the proteins identified in the pericarp proteome of seeds from two popcorn inbred lines (P11 and P16) in two developmental stages (15 and 25 DAP) were intrinsically involved in the key processes associated with pericarp development and thickening (Fig 1, S1 Table). Metabolic process was the category that presented the highest number of associated proteins (29.3%), emphasizing the primary metabolic process with 66% of this total. The molecular functions of catalytic activity and binding had 33% and 25%, respectively. For cellular components, the categories of cell (13.8%) and cell part (13.8%) were the ones with the highest number of associated proteins. These results are very similar to those obtained from the pericarp of the N04 popcorn inbred line evaluated in three stages of seed development (10, 20, and 33 DAP) [29]. The only difference observed was an inversion between the molecular functions binding and catalytic activity, with 45.87% and 41.46% of associated proteins, respectively [29].

Ten proteins that regulate the organization of cell walls aiding in assembly, arrangement of constituent parts, or disassembly of cell walls were identified (Fig 1). Among them, expansin (Q94KT7) promotes the loosening and extension of cell walls, interrupting the non-covalent bond between cellulose microfibrils and matrix glucans [30,31]. Furthermore, this protein appears to be strongly bound to xylans present in type II primary cell walls found in grasses such as popcorn maize [31].

The identification of proteins that regulate the organization of cell walls is explained by the processes involved in pericarp development. The pericarp is a maternal tissue originating from the ovary walls. After undergoing a period of cellular expansion in the first 9 DAP, it collapses between 10 and 18 DAP. This phenomenon occurs after partial resorption of cells to form the inner and outer pericarp, adjusting to the expanding endosperm [32]. The 15 DAP time point analyzed in the present work comprises this period of cellular collapse and reorganization.

Further in the development process, at approximately 20 DAP, the cells of the inner pericarp collapse, while those of the outer pericarp elongate, causing a significant thickening of the cell walls. Consequently, the pericarp thickens [32], forming a strong protective layer. This thickening continues gradually until the later stages of maturation [25]. The 25 DAP time point evaluated in the present study is included in this thickening phase. Therefore, pericarp reorganization, expansion, and thickening processes, which occur between 10 and 20 DAP, directly affect the number of identified proteins and the complexity of observed biological and biochemical processes. Pericarp thickness is a complex trait that has high heritability in the narrow sense (55-82%), involving additive, dominant, and epistatic effects [27, 33], with QTLs and candidate genes associated with it [24, 35].

### Differentially abundant proteins

The protein profile, especially the differentially abundant ones, contributed greatly to understanding the gene regulation during kernel pericarp development in the two inbred lines of popcorn analyzed. It was possible to observe that the lines showed regulation differences even at the same point in the plant cycle.

The highest number of up-accumulated proteins was identified in the inbred line P16 (low expansion capacity), 83% at 25 DAP, all allocated to global metabolic pathways (Fig 2, S3 Table). The remaining proteins were associated with secondary metabolism. The same behavior was observed in the inbred line with high expansion capacity (P11); 77% of the proteins were allocated to global metabolic pathways.

The number of up-accumulated proteins identified in the two popcorn inbred lines and their metabolic pathways was lower and different from what was observed in the pericarp of N04 [29]. This difference is probably due to genotype, method of analysis, and developmental stages. The authors pointed out that the number of proteins identified varies according to the stage of development evaluated.

### Candidate proteins

Three differentially abundant proteins stood out in the two popcorn inbred lines, two in P16 (low expandability) and one in P11 (high expandability). Two are peroxidases, and one is a xyloglucan endotransglucosylase/hydrolase (XTH). Peroxidases participate in phenylpropanoid biosynthesis, specifically in lignin biosynthesis, and xyloglucan endotransglucosylase/hydrolase is an enzyme that promotes the cleavage and reassembly of xyloglucan molecules [31].

### Peroxidases

The final step of lignin biosynthesis is the polymerization of monolignols (sinapyl alcohol, S unit; coniferyl alcohol, G unit; and ρ-coumaryl alcohol, H unit) in the secondary cell wall domains by peroxidases and laccases [14–17]. The function of these enzymes is conditioned by spatio-temporal regulation of gene expression, as well as their differentiated oxidation [14]. In this study, the peroxidases B6T6D4 and A0A1D6E530 were up-accumulated in P16/15 DAP and P11/25 DAP, respectively.

As previously mentioned, 25 DAP is the developmental stage in which the secondary cell wall of the pericarp thickens, making it more rigid, that is, more lignified. Therefore, the greater accumulation of the peroxidase A0A1D6E530 in the pericarp of P11 makes its cell walls more rigid. Thus, it indicates why the pericarp of this inbred line resists longer to increased pressure inside the kernel during the popping process [6, 7]. The content and composition of lignin present in the pericarp of three popcorn inbred lines, including the two analyzed in the present work, have been evaluated [18]. According to the authors, there was a statistically significant difference (5% probability by the F test) between the lignin content present in the pericarp of the popcorn inbred lines. The lines with higher expansion volume values, including P11, presented higher lignin contents, concluding that lignin positively contributes to the expansion volume [18].

The gene that encodes the peroxidase A0A1D6E530 is *Zm00001eb076190*, also called *Sb03g0468102*, located on chromosome 2 (26,292,971-26,294,428) and available in the Gramene database (https://www.gramene.org/). This gene was shown to be up-regulated during 25 DAP in the highly expandable line P11.

P16 (low expansion capacity) presented increased peroxidase (B6T6D4) accumulation at 15 DAP. This stage is inserted in the period of cellular collapse and formation of the internal and external pericarp, with an adjustment for endosperm expansion [32]. Class III peroxidases, where A0A1D6E530 and B6T6D4 belong, present spatio-temporal regulation of gene expression, as well as differentiated oxidation properties [14]. Therefore, the regulation of peroxidase B6T6D4 in P16 was probably directed at the initial events of pericarp collapse and reorganization. Between 15 and 18 DAP, the pericarp reaches its maximum thickness in the seed basal region [36], thus indicating that the peroxidase B6T6D4 from P16 is involved in pericarp thickening. In contrast, the peroxidase A0A1D6E530 is linked to lignin deposition, making the pericarp of P11 (high expansion capacity) more resistant.

Comparing the results obtained in the present work with those available in the literature, especially with the 11 MetaQTLs associated with increased popcorn expansion volume [35], it can be seen that the gene Zm00001eb076190 that encodes a peroxidase (A0A1D6E530) had not yet been reported. Therefore, this gene is an important addition to understanding the studied trait and can be used for introgression in breeding programs aiming to improve popcorn expansion capacity.

### Xyloglucan endotransglucosylase/hydrolase

In P16 (low expansion capacity), at both 15 and 25 DAP, xyloglucan endotransglucosylase/hydrolase (XTH) was up-accumulated. This protein is a member of the GH16 (glycosidic hydrolase) family, actively acting to break β-1,4 or β-1,3 glycosidic bonds in various CAZyme glucans and galactans (Carbohydrate-Active Enzymes Database) [11, 37]. This enzyme exhibits transglycosylase (XET) and glycosidic hydrolase (XEH) activity [31]. Like expansins, xyloglucan endotransglucosylase/hydrolase is versatile in the metabolic process of cell wall biogenesis and organization [31, 38–40]. When XET acts on xyloglucan, it provides a reversible cell wall loosening mechanism necessary for plant cell expansion [41], while XEH causes irreversible extension [11]. In addition, transglycolysation predominates over hydrolysis unless the receptor concentration is very low [42].

Some XTH gene families have already been identified. For instance, *Arabidopsis thaliana* presents 33 [43] genes while tomato *(Solanum lycopersicum* L.) has 25 members [44]. In monocots, such as wheat [45] and rice [46], 57 and 29 genes were identified, respectively. In maize, the number of genes belonging to the XTH family is unknown. However, *Zm1005* was identified [47]. This gene was validated on March 6, 2019, and published in the National Center for Biotechnology Information, receiving the symbol *LOC542059* (or *xth1* and *GRMZM2G119783*), locus tag *ZEAMMB73_Zm00001d024386*, located on chromosome 10, and containing 1,475 nucleotides (68,324,472 to 68,325.946) [48].

The expression of XTH family genes is normally tissue-specific and developmental stage-dependent, as observed in *Oryza sativa* [43], *Triticum durum* [49], and *Hordeum vulgare* [40]. XTHs act on xyloglucan, which is not the main hemicellulose that contributes to cell wall organization in *Z. mays*. Nevertheless, it is suggested that even though it is less abundant, xyloglucan metabolism plays a central role in restructuring the cell wall of commelinoid monocots, such as maize [10]. In addition, hydrolases also play a role in cell extension [31].

Considering the above, it would be possible to hypothesize that xyloglucan endotransglucosylase/hydrolase (B4FHS5) affects the pericarp cell walls of P16 (low expansion capacity) since this protein was up-accumulated in both developmental stages, indicating that the *GRMZM2G119783* gene was up-regulated. In this context, it is suggested that xyloglucan endotransglucosylase/hydrolase participates in regulating the metabolic process of biogenesis and cell wall organization together with expansins, promoting the irreversible loosening of the pericarp cell walls of P16 (low expansion capacity). Thus, it makes it thinner and compromises its integrity, making it unable to withstand the pressure generated inside the kernel. In this way, the pericarp is ruptured before the endosperm is completely gelatinized, causing P16 to have a lower expansion capacity. However, this hypothesis needs to be further investigated.

## Conclusion

Our data strongly suggest that the peroxidase A0A1D6E530 increases the expansion capacity of popcorn kernels by increasing the lignin content. On the other hand, the xyloglucan endotransglucosylase/hydrolase B4FHS5 has been shown to play an important role, yet to be elucidated, in the decrease of expandability. Since expansion capacity is a polygenic trait, these two proteins and their respective genes stand out as important candidates in this pathway, contributing positively and negatively to the popping process, respectively. This information is unprecedented and can be used by popcorn breeding programs.

## Supporting information

**S1 Fig.** Seeds of inbred lines P16 (expansion volume 14 mL g^-1^) and P11 (expansion volume 30 mL g^-1^) in both developmental stages.

(TIFF)

**S2 Fig.** Processing of seeds for further protein extraction. a) Ear of inbred line P11 (15 DAP); b) removal of individual seeds from the ear with a scalpel; c) longitudinal cut performed in the kernel pericarp; d) pericarps separated from the seeds and frozen. (TIFF)

**S3 Fig.** Protein profiles obtained in the pericarp of popcorn seeds of lines P11 and P16 in the four treatments analyzed, namely, T1 (P11/15 DAP), T2 (P16/15 DAP), T3 (P11/25 DAP), and T4 (P16/25 DAP).

(TIFF)

**S1 Table.** Specific proteins identified in the pericarp of inbred lines P11 and P16.

(XLSX)

**S2 Table.** GO terms referring to biological processes (P), molecular function (F), and cellular components (C) for pericarp proteins.

(XLSX)

**S3 Table.** Differentially abundant proteins identified in the pericarp of P11 and P16 inbred lines, considering the RI (P11_15DAp_/P16_15DAP_) and RII (P112_5DAP_/P162_5DAP_) ratios. (XLSX)

**S4 Table.** Metabolic pathways that showed up-accumulated proteins in the pericarp of popcorn inbred lines P11 and P16 during seed development at 15 DAP (ratio RI) and 25 DAP (ratio RII).

(XLSX)

## Acknowledgments

The authors would like to thank the Coordination for the Improvement of Higher Education Personnel (Coordenação de Aperfeiçoamento de Pessoal de Nível Superior – CAPES), Prof. Carlos Alberto Scapim for providing the seeds of the popcorn maize lines, and the Proteomics Laboratory (LABPROT), Institute of Chemistry, Federal University of Rio de Janeiro (UFRJ) for technical support in the proteome analysis.

## Author Contributions

### Conceptualization

Adriana Gonela, Isaac Romani, Gilberto Barbosa Domont, Fábio César Sousa Noqueira.

### Data curation

Talita Mayara de Campos Jumes Gemelli, Isaac Romani, Natália Ferreira dos Santos.

### Formal analysis

Talita Mayara de Campos Jumes Gemelli, Isaac Romani, Natália Ferreira dos Santos, Gilberto Barbosa Domont, Fábio César Sousa Noqueira.

### Funding Acquisition

Adriana Gonela, Gilberto Barbosa Domont, Fábio César Sousa Noqueira.

### Investigation

Talita Mayara de Campos Jumes Gemelli, Isaac Romani, Natália Ferreira dos Santos.

### Methodology

Adriana Gonela, Carlos Alberto Scapim, Gilberto Barbosa Domont, Fábio César Sousa Noqueira, Talita Mayara de Campos Jumes Gemelli, Isaac Romani, Natália Ferreira dos Santos.

### Project administration

Adriana Gonela, Gilberto Barbosa Domont, Fábio César Sousa Noqueira, Talita Mayara de Campos Jumes Gemelli, Isaac Romani, Natália Ferreira dos Santos.

### Resources

Adriana Gonela, Carlos Alberto Scapim, Talita Mayara de Campos Jumes Gemelli, Isaac Romani, Natália Ferreira dos Santos.

### Software

Talita Mayara de Campos Jumes Gemelli, Isaac Romani, Natália Ferreira dos Santos.

### Supervision

Adriana Gonela, Gilberto Barbosa Domont, Fábio César Sousa Noqueira, Maria de Fátima P.S. Machado.

### Validation

Adriana Gonela, Carlos Alberto Scapim, Gilberto Barbosa Domont, Fábio César Sousa Noqueira, Maria de Fátima P.S. Machado.

### Visualization

Adriana Gonela, Carlos Alberto Scapim, Gilberto Barbosa Domont, Fábio César Sousa Noqueira, Maria de Fátima P.S. Machado, Talita Mayara de Campos Jumes Gemelli, Isaac Romani, Natália Ferreira dos Santos.

### Writing – original draft, Writing – review & editing

Adriana Gonela, Talita Mayara de Campos Jumes Gemelli, Isaac Romani, Natália Ferreira dos Santos.

## References

1. Sweley JC, Rose DJ, Jackson DS. Quality traits and popping performance considerations for popcorn (*Zea mays* Everta). Food Reviews International. 2013; 29(2):157–177.

2. Mishra G, Joshib DC, Pandaa BK. Popping and puffing of cereal grains: a review. Journal of Grain Processing and Storage. 2014; 1(2):34–46.

3. Park D, Maga JA. Effects of storage temperature and kernel physical condition on popping qualities of popcorn hybrids. Cereal Chemistry. 2000; 79(4):572–575.

4. Ziegler KE. Popcorn. In: Halluer AR, editor. Specialty corns. CRC press: USA; 2001. p. 199–234.

5. Borras F, Seetharaman K, Yao N, Robute JL, Percibaldi NM, Eyherabide GH. Relationship between popcorn composition and expansion volume and discrimination of corn types by using zein properties. Cereal Chemistry. 2006; 83(1):86–92.

6. Silva WJ, Vidal BC, Pereira AC, Zerbeto M, Vargas H. What makes popcorn pop. Nature. 1993; 362: 417–417.

7. Hoseney RC, Zeleznak K, Abdelrahman A. Mechanism of popcorn popping. Journal of Cereal Science. 1983; 1(1):43–52.

8. Scheller HV, Ulvskov P. Hemicelluloses. Annual Review of Plant Biology. 2010; 61:263–289.

9. Carpita NC, Gibeaut DM. Structural models of primary cell walls in flowering plants: consistency of molecular structure with the physical properties of the cell wall during growth. The Plant Journal. 1993; 3(1):1–30.

10. Yokoyama R, Rose JKC, Nishitani K. A surprising diversity and abundance of xyloglucan endotransglucosylase/hydrolases in rice: classification and expression analysis. Plant Physiology. 2004; 134(3):1088–1099.

11. Eklöf JM, Brumer H. The XTH gene family: an update on enzyme structure, function, and phylogeny in xyloglucan remodeling. Plant Physiology. 2010; 153(2):456–66.

12. Moreira-Vilar FC, Siqueira-Soares RC, Finger-Teixeira A, Oliveira DM, Ferro AP, Rocha GJ, et al. The acetyl bromide method is faster, simpler and presents better recovery of lignin in different herbaceous tissues than Klason and thioglycolic acid methods. PLoS One. 2014; 9, e110000.

13. Ithal N, Recknor J, Nettleton D, Maier T, Baum TJ, Mitchum MG. Developmental transcript profiling of cyst nematode feeding cells in soybean roots. Mol Plant Microbe Interact. 2007; 20, 510–525.

14. Shigeto J, Tsutsumi Y. Diverse functions and reactions of class III peroxidases. New Phytologist. 2016; 209: 1395–1402. doi: 10.1111/nph.13738.

15. Laitinen T, Morreel K, Delhomme N, Gauthier A, Schiffthaler B, Nickolov K, et al. A key role for apoplastic H2O2 in Norway spruce phenolic metabolism. Plant Physiol. 2017; 174:1449–1475.

16. Dixon RA, Barros J. Lignin biosynthesis: old roads revisited and new roads explored. Open Biol. 2019; 9: 190215. http://dx.doi.org/10.1098/rsob.190215.

17. Tobimatsu Y, Schuetz M. Lignin polymerization: how do plants manage the chemistry so well? Current Opinion in Biotechnology. 2019; 56:75–81.

18. Damasceno Junior C, Godoy S, Gonela A, Scapim CA, Grandis G, Santos WD, et al. Biochemical composition of the pericarp cell wall of popcorn inbred lines with different popping expansion. Current Research in Food Science. 2022; 5:102–106.

19. Robbins ML, Roy A, Wang PH, Gaffoor I, Sekhon RS, de Buanafina MM. Comparative proteomics analysis by DIGE and iTRAQ provides insight into the regulation of phenylpropanoids in maize. Journal of Proteomics. 2013; 93:254–275.

20. Thingholm TE, Palmisano G, Kjeldsen F, Larsen MR. Undesirable charge-enhancement of isobaric tagged phosphopeptides leads to reduced identification efficiency. J Proteome Res. 2010; 9:4045–52.

21. Nogueira FCS, Palmisano G, Schwämmle V, Campos FAP, Larsen MR, Domont GB, et al. Performance of isobaric and isotopic labeling in quantitative plant proteomics. J Proteome Res. 2012; 11:3046–52.

22. Santos MDM, Lima DB, Fischer JSG, Clasen MA, Kurt LU, Camillo-Andrade AC, et al. Simple, efficient and thorough shotgun proteomic analysis with PatternLab V. Nature protocols. 2022. https://doi.org/10.1038/s41596-022-00690-x

23. Kanehisa M, Goto S, Sato Y, Furumichi M, Tanabe M. KEGG for integration and interpretation of large-scale molecular data sets. Nucleic Acids Research. 2015; 10(1):109–114.

24. Tian T, Liu Y, Yan H, You Q, Yi X, Du Z, et al. AgriGO v. 2.0: a GO analysis toolkit for the agricultural Community. Nucleic Acids Res. 2017; 45:W122–W129.

25. Richardson DL. Pericarp thickness in popcorn. Agronomy Journal. 1960; 52(2):77–80.

26. Mohamed AA, Ashman RB, Kirleis AW. Pericarp thickness and other kernel physical characteristics relate to microwave popping quality of popcorn. Journal of Food Science. 1993; 58 (2):342–346.

27. Ho LC, Kannenberg LW, Hunter RB. Inheritance of pericarp thickness in short season maize inbreds. Canadian Journal of Genetics and Cytology. 1975; 17(4):621–629.

28. Tandjung AS, Janaswamy S, Chandrasekaran R, Aboubacar A, Hamaker BR. Role of the pericarp cellulose matrix as a moisture barrier in microwaveable popcorn. Biomacromolecules. 2005; 6(3):1654–1660.

29. Dong Y, Wang Q, Zhang L, Du C, Xiong W, Chen X, et al. Dynamic proteomic characteristics and network integration revealing key proteins for two kernel tissue developments in popcorn. PLoS One. 2015; 10(11):77–86.

30. Sampedro J, Cosgrove DJ. The expansin superfamily. Genome Biology. 2005; 6:242. doi:10.1186/gb-2005-6-12-242.

31. Carpita NC, Ralph J, McCann MC. The cell wall. In: Buchanan BB, Gruissem W, Jone RL, editors. Biochemistry & Molecular Biology of Plants. Oxford: John Wiley & Sons, Ltd.; 2015. p. 45–110.

32. García-Lara S, Bergvinson DJ, Burt AJ, Ramputh AI, Díaz-Pontones DM, Arnason JT. The role of pericarp cell wall components in maize weevil resistance. Crop Science. 2004; 44:1546–1552.

33. Ito GM, Brewbaker JL. Genetic analysis of pericarp thickness in progenies of eight corn hybrids. J Amer Soc Hort Sci. 1991; 116(6):1072–1077.

34. Wu X, Wang B, Xie F, Zhang L, Gong J, Zhu W, et al. QTL mapping and transcriptome analysis identify candidate genes regulating pericarp thickness in sweet corn. BMC Plant Biology. 2020; 20:117. doi.org/10.1186/s12870-020-2295–8.

35. Kaur S, Rakshit S, Choudhary M, Das AK, Kumar RR. Meta-analysis of QTLs associated with popping traits in maize (*Zea mays* L.). PLoS One. 2021; 16(8): e0256389. https://doi.org/10.1371/journal.pone.0256389.

36. Randolph LF. Developmental morphology of the caryopsis in maize. J Agr Res. 1936; 53:881–916.

37. Rose JK, Braam J, Fry SC, Nishitani K. The XTH family of enzymes involved in xyloglucan endotransglucosylation and endohydrolysis: current perspectives and a new unifying nomenclature. Plant Cell Physiology. 2002; 43(12):1421–1435.

38. Hara Y, Yokoyama R, Osakabe K, Toki S, Nishitani K. Function of xyloglucan endotransglucosylase/hydrolases in rice. Annals of Botany. 2014; 114(6):1309–1318.

39. Sharples SC, Nguyen-Phan TC, Fry SC. Xyloglucan endotransglucosylase/hydrolases (XTHs) are inactivated by binding to glass and cellulosic surfaces, and released in active form by a heat-stable polymer from cauliflower florets. Journal of Plant Physiology. 2017; 218:135–143.

40. Fu, MM, Liu C, Wu F. Genome-wide identification, characterization and expression analysis of xyloglucan endotransglucosylase/hydrolase genes family in barley *(Hordeum vulgare)* Molecules. 2019; 24:1935–1946.

41. Fry SC, Smith RC, Renwick KF, Martin DJ, Hodge SK, Matthews, KJ. Xyloglucan endotransglycosylase, a new wall-loosening enzyme activity from plants. Biochemical Journal. 1992; 282(pt 3):821–828.

42. Fanutti C, Gidley MJ, Reid JS. Action of a pure xyloglucan endo-transglycosylase (formerly called xyloglucan-specific endo-(1→4)-beta-D-glucanase) from the cotyledons of germinated nasturtium seeds. The Plant Journal. 1993; 3(5):691–700.

43. Yokoyama R, Nishitani K. A comprehensive expression analysis of all members of a gene family encoding cell-wall enzymes allowed us to predict cis-regulatory regions involved in cell-wall construction in specific organs of Arabidopsis. Plant Cell Physiology. 2001; 42(10):1025–1033.

44. Saladié M, Rose JK, Cosgrove DJ, Catalá C. Characterization of a new xyloglucan endotransglucosylase/hydrolase (XTH) from ripening tomato fruit and implications for the diverse modes of enzymatic action. Plant Journal. 2006; 47(2): 282–295.

45. Liu Y, Liu D, Zhang H, Gao H, Guo X, Wang D, et al. The α-and β-expansin and xyloglucan endotransglucosylase/hydrolase gene families of wheat: Molecular cloning, gene expression, and EST data mining. Genomics. 2007; 90(4):516–529.

46. Uozu S, Tanaka-Ueguchi M, Kitano H, Hattori K, Matsuoka M. Characterization of XET related genes in rice. Plant Physiology. 2000; 122(3): 853–859.

47. Saab IN, Sachs MM. A flooding-induced xyloglucan endo-transglycosylase homolog in maize is responsive to ethylene and associated with aerenchyma. Plant Physiology. 1996; 112(1): 385–391.

48. NCBI. LOC542059 xyloglucan endo-transglycosylase/hydrolase [Zea mays]. Available from: https://www.ncbi.nlm.nih.gov/gene?Db=gene&Cmd=DetailsSearch&Term=542059.

49. Iurlaro A, Caroli MD, Sabella E, Pascali MD, Rampino P, Bellis LD, et al. Drought and heat differentially affect XTH expression and XET activity and action in 3-day-old seedlings of durum wheat cultivars with different stress susceptibility. Frontiers in Plant Science. 2016; 7:10–26.

